# Automatic Feedback Control for Resource-Aware Characterisation of Genetic Circuits

**DOI:** 10.1101/2025.04.01.646303

**Authors:** Kirill Sechkar, Harrison Steel

**Affiliations:** Department of Engineering Science, University of Oxford, Oxford, OX1 3PJ, UK

**Keywords:** Synthetic Biology, Genetic Expression, Bifurcation, Feedback, Automatic Control, Mathematical Models

## Abstract

Many applications of engineered cells are enabled by genetic circuits – networks of genes regulating each other to process signals. Complex circuits are built by combining standardised modular components with different functions. Nonetheless, genes in a cell compete for the same limited pool of cellular resources, causing unintended interactions that violate modularity. Thus, circuit components behave differently when combined versus when observed in isolation, which can compromise a biotechnology’s predictability and reliability. To forecast steady-state interactions between modules, experimental protocols for characterising their resource competition properties have been proposed. However, they rely on open-loop batch culture techniques in which dynamic control signals cannot be applied to cells. Consequently, these experimental methods have limited predictive power, as they may fail to capture all possible steady states, such as repelling equilibria that would not be approached by a system without external forcing. In contrast, we propose a novel, comprehensive protocol for characterising the resource-dependence of genetic modules’ performance. Based on the control-based continuation technique, it captures both stable and unstable steady states by applying stabilising cybergenetic feedback with an automated cell culturing platform. Using several models with different degrees of complexity, we simulate applying our pipeline to a self-activating genetic switch. This case study illustrates how informative characterisation of a genetic module with automatic feedback control enables reliable forecasting of its performance when combined with any other circuit component. Hence, our protocol promises to restore predictability to the design of genetic circuits from standardised components.

## I. INTRODUCTION

### A. Challenges of resource-aware circuit characterisation

Engineered cells’ biosensing, biomanufacturing and therapeutic applications are powered by gene circuits — networks of genes that regulate each other to perceive, process and react to complex stimuli [1], [2]. Engineering principles of modularity and standardisation allow to construct increasingly sophisticated circuits by combining genetic modules with different functions [3]. However, all genes in a cell compete for the same limited pool of expression resources (such as ribosomes), which introduces unintended indirect interactions between circuit components. Due to these resource couplings, genetic modules behave differently when characterised in isolation and when joined into a circuit, which can compromise the predictability and reliability of biotechnologies [2], [4].

This predictability challenge has spawned resource-aware mathematical models of gene expression incorporating resource coupling dynamics into their ordinary differential equations (ODEs) [1], [2], [5], [6]. However, modelling requires knowing a system’s parameters, whose estimation from experimental data may be uncertain, expensive and time-consuming [7]. Instead of inferring all parameters, one can measure lumped factors capturing a genetic module’s empirically observed input-output relationships, which are reproduced by many different modelling frameworks [1], [4]. This allows to forecast competition between any two genetic modules for which these quantities are known. For instance, McBride and Del Vecchio suggested predicting steady-state circuit behaviours based on ‘resource demand’ factors, measured for each genetic module of interest by observing cells which express it alongside a constitutive reporter gene with known resource competition properties (Fig. 1a) [4].

**Fig. 1.**
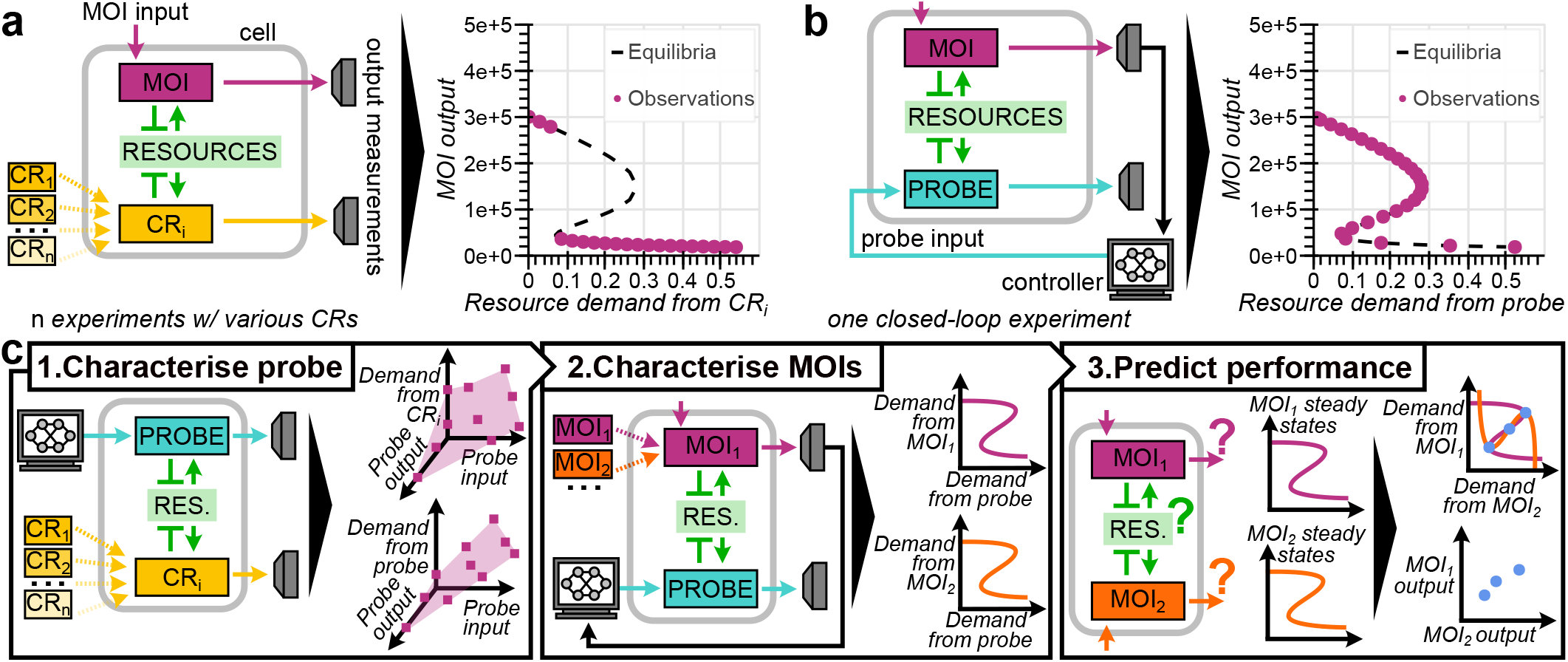
Overview of the proposed method. (a) Conventional resource competition characterisation, in which a genetic module of interest MOI is co-expressed with different constitutive reporters CR_1_, CR_2_, … CR_n_ one-by-one, may not capture all steady states of the system. (b) CBC characterisation, where the MOI’s output is steered to different reference values by controlling resource competition in the cell via a co-expressed probe module, captures equilibria all along the bifurcation curve. Observations in (a–b) are actual simulations of the model in Section II, highlighting the extant method’s limitations compared to CBC. (c) Steps of the proposed protocol for characterising MOIs with CBC and predicting their steady-state outputs in the light of resource competition.

However, such open-loop experimental methods may only capture one among many possible equilibria of a system (Fig. 1a) [7], [8]. At the same time, multistability of gene circuits underlies cellular development and adaptability to environmental changes in nature [3], [9] and is often observed in synthetic biology [3], [7], [10]. Competition may not only shift a circuit’s equilibria [2], [4], but also cause bifurcations that qualitatively alter their number and identity [3], [11]. These bifurcations, for instance, have been leveraged by a biomolecular controller improving biotechnologies’ robustness to mutations [12]. Meanwhile, a system’s unstable steady states can be hard to capture in open-loop setups – as systems diverge from them without forcing – yet be vital for predicting circuit behaviour, since they delimit the basins of attraction of different stable equilibria (e.g. representing different states of a memory device) [3], [11]. Hence, reliably forecasting genetic modules’ performance requires observing their multistability and bifurcations associated with resource competition, which extant experimental protocols fail to achieve.

### B. Circuit characterisation with feedback control

This paper outlines a novel experimental approach aimed at addressing the current limitations of resource competition characterisation methods for genetic modules. Instead of subjecting a module of interest to constant resource demands created by constitutive reporters, we leverage a ‘probe’ module, the expression of whose genes is dynamically steered by a feedback control signal from an automated cell culture platform. This enables control-based continuation (CBC), an emerging model-free testing algorithm recently extended to biology [7]. In CBC, stabilising feedback steers the system towards a steady state – even if it is normally unstable – allowing to observe it experimentally. Driving the plant through several equilibria in sequence produces a bifurcation diagram revealing all of the system’s steady states at different extents of resource competition (Fig. 1b) [8], [13].

Our protocol for characterising genetic modules and predicting their performance involves three steps (Fig. 1c). A probe module with known competition properties is established. Then, a genetic module of interest (MOI) is characterised using this probe in a CBC setup. Finally, bifurcation diagrams for different modules are combined to predict their behaviour in the same host cell as they compete for its resources.

As an example of applying this approach, this paper considers a self-activating genetic switch module, described using a basic resource-aware ODE model outlined in Section II and picked because its qualitatively resource-dependent steady-state behaviour has been observed experimentally [3] and leveraged in circuit design [12]. In Section III, all three steps of our protocol are performed *in silico*. In Section IV we showcase our method’s model-agnosticism by repeating it for another, more complex resource-aware model of an engineered *E. coli* cell. Our method’s implementability *in vivo*, potential limitations and applications are discussed in Section V.

## II. MODEL

### A. Gene expression ODEs

Most simulations and derivations in this paper are performed for a basic model of an *E. coli* cell expressing its native genes and some synthetic genes added to it. Heavily simplified to clearly illustrate our protocol, this framework still incorporates key resource competition considerations of finite protein expression resources (most crucially for bacteria, ribosomes [6]) and constraints on the cell’s total protein mass [1], [2], [14]. The cell’s native genes are lumped into two coarse-grained classes, each treated as a single gene – ribosomes *r*, which enable protein synthesis, and all other genes *o*. Together with the synthetic genes, *r* and *o* comprise the set *S* of all genes being expressed. The concentration of gene *i* ∈ *S* proteins, denoted as *p*_*i*_, has its dynamics described by (1):

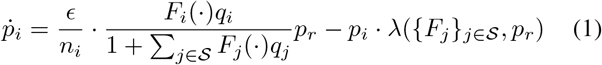

Here, protein production is a single-step reaction with rate *ϵ* in amino acids per ribosome per hour, hence the need to divide it by the protein’s length in amino acids *n*_*i*_. The factor before the total abundance of ribosomes *p*_*r*_ captures the fraction of resources allocated towards making protein *i*, where *q*_*i*_ is its maximum possible resource demand and 0 *≤F*_*i*_(*·*) *≤* 1 is the regulation function with gene-specific form and arguments. Native genes are treated as constitutive (hence *F*_*r*_ *≡ F*_*o*_ *≡* 1) because their regulation is assumed unchanging in a given fixed culture condition, but this simplification is lifted in Section IV. Protein removal predominantly occurs by dilution as the cell grows and expands in volume at rate λ [1]. Protein mass density in the cell *M* is constant [14], so this growth rate must equal the total rate of protein synthesis, i.e.:

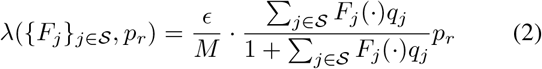

A fluorescent protein must maturate before it can be measured [15]. For such genes, we separately consider nascent and mature protein concentrations 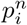 and 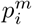; only the latter variable can be experimentally observed by an instrument (e.g. a microscope). The split ODE (1) becomes ODEs (3a)–(3b), where *µ*_*i*_ is the maturation rate (note that 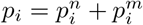):

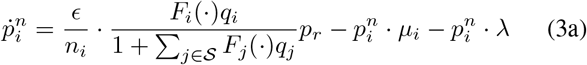

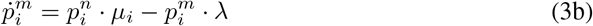

### B. Genetic modules

Besides the cell’s native genes, in this paper we consider three kinds of synthetic gene circuit modules, shown in Fig. 2:

**Fig. 2.**
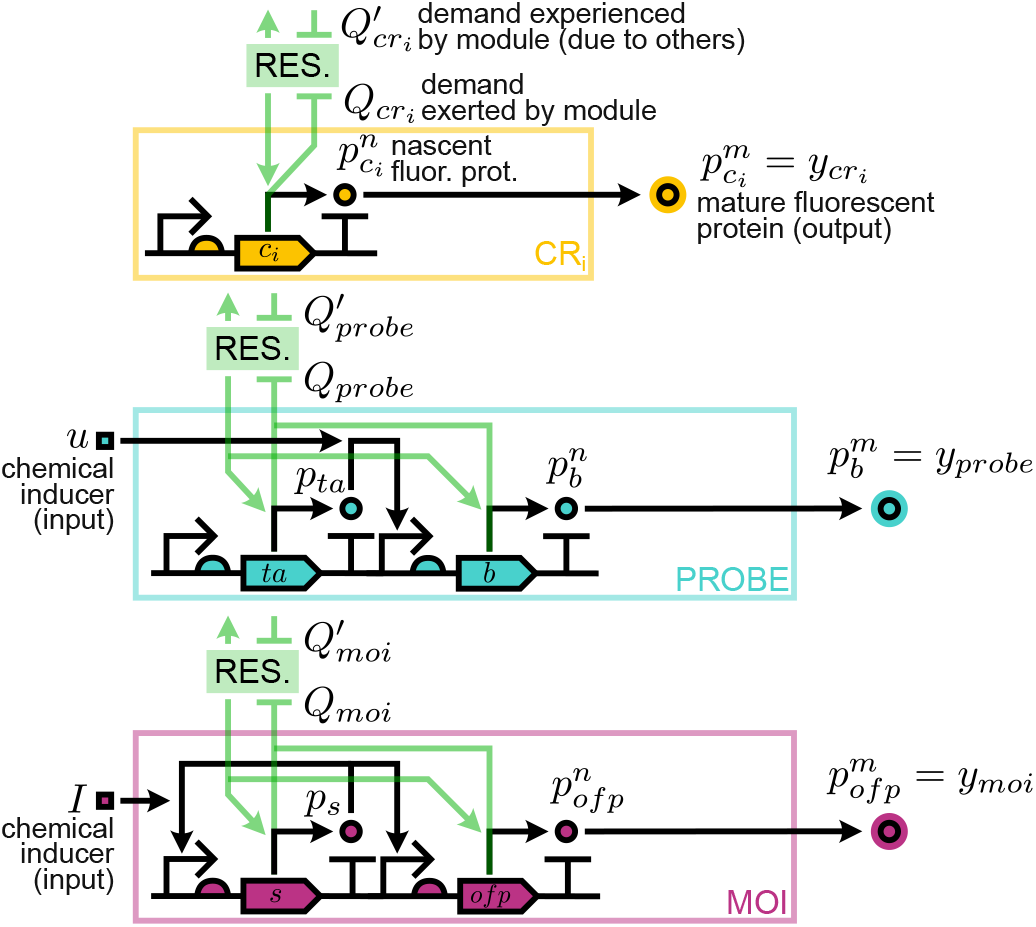
Genetic modules considered in this study and described in Section II-B

1. <u>Constitutive reporters</u>, used in conventional characterisation experiments. Each reporter is an unregulated fluorescent protein gene *c*_*i*_. Hence, its regulation function is 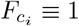, and its protein dynamics are given by ODEs (3a)–(3b).
2. <u>Probe module</u>, steered by the controller to manipulate the resource demand in the cell, e.g. by changing the concentration of a chemical inducer *u* in the culture medium. This modulates the activity of constitutive transcription activator (*ta*) proteins, which must be bound by an inducer in order to upregulate the expression of a fluorescent protein *b*. The *ta* gene’s expression is governed by ODE (1) and that of *b* by ODEs (3a)–(3b). Whilst *F*_*ta*_ *≡*1, *F*_*b*_ is given by (4), which captures inducer-activator protein binding with a half-saturation constant *K*_*u*_, as well as cooperative binding between the inducer-activator complexes and the *b* gene’s DNA with a half-saturation constant *K*_*b*_ and a Hill coefficient *η*_*b*_. Without activators, gene *b* has low baseline expression *F*_*b*,0_.

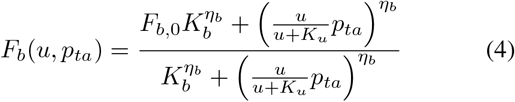
3. <u>Module of interest (self-activating genetic switch)</u>, comprising a switch gene *s* and an output fluorescent protein gene *ofp*, whose protein dynamics are respectively given by ODE (1) and ODEs (3a)–(3b). Like *ta, s* is a transcription activator. However, it is bound by a different inducer with unchanging concentration throughout the experiment, so the fraction of inducer-bound switch proteins is a constant *I*. Being self-activating, the switch upregulates both its own and protein *ofp*’s synthesis. We thus have (5) with *K*_*s*_, *η*_*s*_ and *F*_*s*,0_ analogous to the corresponding parameters in (4):

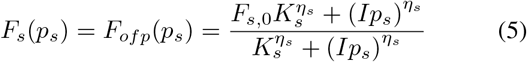

### C. Notation for module resource demands and outputs

Let the cell host two genetic modules with gene sets ℳ_1_ and ℳ_2_, so *S* = {*r, o*} ∪ ℳ_1_ ∪ ℳ_2_. We normalise the synthetic genes’ steady-state resource demands to those of the native genes as per (6), where the bar indicates equilibrium values:

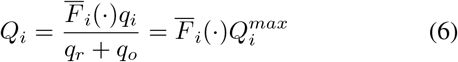

These normalised factors are summed to find for each module the total resource demand that its genes *exert* on the cell and the total demand it *experiences* due to the other module. For the first module, these quantities are respectively 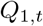 and 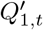 in (7). The demand exerted by one module is experienced by the other, so for the second module we have 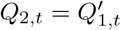 and 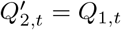.

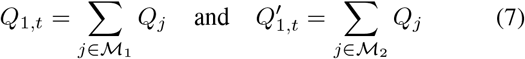

Consider the steady-state observations made for cells expressing a given pair of modules (this scenario is denoted as _1+2_). First, there are these modules’ corresponding outputs 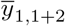 and 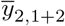, i.e. the levels of mature output fluorescent proteins, whose emission is observed by a microscope. Second, the cell growth rate 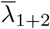 is measurable as the rate of change of cell length in microscopy images [10]. A module can also be observed alone in the cell, without any competitors (save for the native genes) – e.g. for module 1, this yields the output 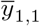 and growth rate 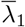. Comparing this to the case of two competing modules and solving (3b) for steady state, the ratio of output protein expression levels can be calculated according to (8), where the output protein maturation rate *µ*_1_ is assumed to be known from literature [15] or prior experiments.

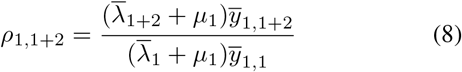

## III. PROTOCOL WALKTHROUGH

### A. Probe characterisation

Since a module of interest’s resource competition properties are originally unknown, in a CBC experiment they must be gauged from inputs and outputs of the probe module co-expressed with it. Importantly, we need to know both the resource demand 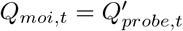 exerted by the MOI on the probe and the demand 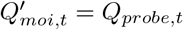 exerted by the probe on the MOI, which may vary if competition affects interactions between the probe’s genes. Whilst the Appendix proves that this can be done unambiguously, the probe’s own resource competition properties must be determined first.

To this end, we establish a library of *n* constitutive reporters by expressing each reporter alone in the cell, then observing pairs of them co-expressed in the same cell. Each reporter module is a single unregulated gene, so their resource demands 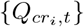 are fixed and can be found as per (9), derived likewise to [4] by considering (8) and (3b) in steady state. For a reporter *cr*_*i*_ observed alone and with another reporter *cr*_*j*_, we have:

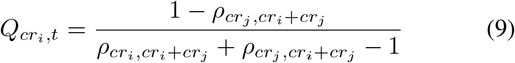

This constitutive reporter library allows to perform an open-loop experiment in which the probe is expressed with each of the reporters, sweeping through *m* inputs and recording fluorescent outputs. As a result, we get *n* × *m* steady-state observations, for which we record 1) probe input 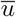, 2) probe output 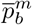, 3) resource demand experienced by the probe 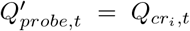 and 4) resource demand exerted by the probe, calculated from the constitutive reporter’s output as:

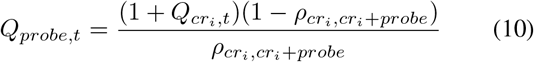

These data points are samples from the mappings 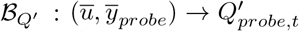 and 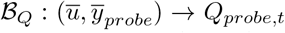 from the probe’s input-output pairs to resource demands experienced and exerted by it. For inputs and/or outputs between the sampled points, 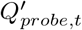 and 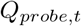 can be interpolated. In CBC characterisation, these mappings will let us gauge resource competition in the cell from probe inputs and outputs.

This procedure is simulated as shown in Fig. 3. For this and all other simulations, we provide model parameters and Python code at https://github.com/KSechkar/CoBaCRAB.

**Fig. 3.**
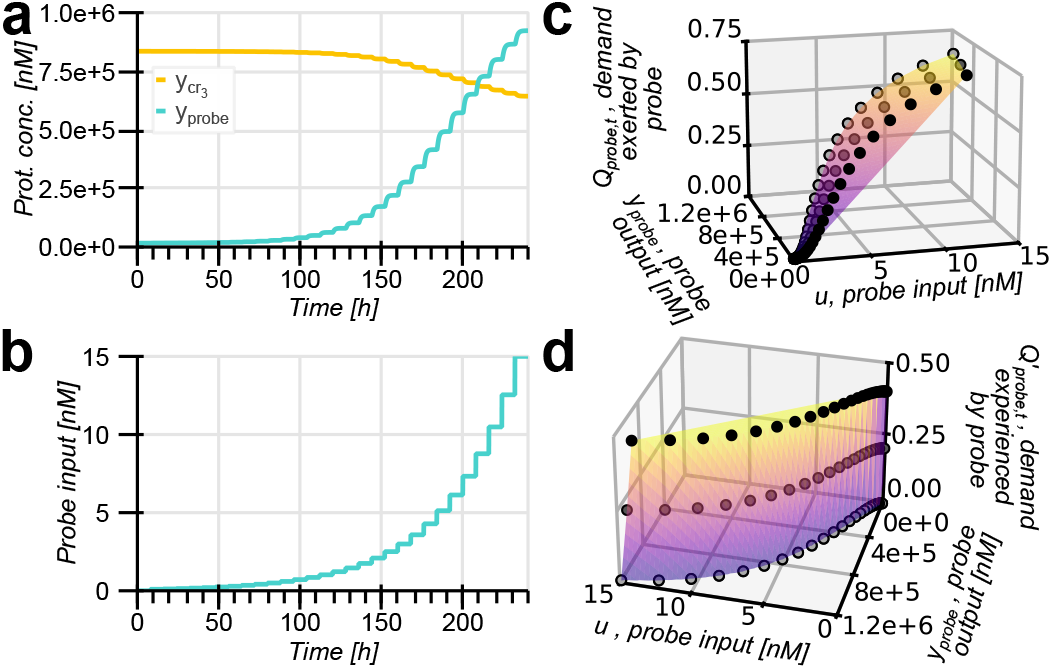
Simulation of probe characterisation with *n* = 3 constitutive reporters and *m* = 16 chemical inputs. (a–b) System trajectory during parameter sweep with the highest-demand reporter *cr*_3_. (c–d) Obtained data points (black) and linear interpolations of the mappings 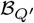 and ℬ_*Q*_ matching the probe’s input-output pairs to the resource demand it experiences and exerts on other modules. Darker points stand for measurements with higher-demand reporters.

### B. Module of interest characterisation with CBC

To characterise a given module of interest, we co-express it with the probe and perform a CBC experiment. The output 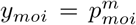 is steered towards a reference *R* by a control signal *u* until both *y*_*moi*_ and *u* reach steady state (Fig. 4a– b), assumed to be achieved after a sufficiently long time or detected by checking slopes of linear curve fits to most recent measurements [7]. Observations at this point correspond to an open-loop system’s steady-state, regardless of whether it was originally stable or unstable (and thus hard to capture). Tracking references *R*_1_, *R*_2_, … *R*_*m*_ one after another provides a comprehensive view of the system’s equilibria [7], [8], [13].

This produces a CBC bifurcation curve, whose points 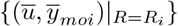 reveal the system’s steady-state input-output combinations. If for each captured steady state we also record probe outputs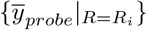, the mappings 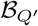 and ℬ_*Q*_ allow to translate the captured bifurcation curve into a resource demand bifurcation curve *C*_*moi*_ whose points

**Fig. 4.**
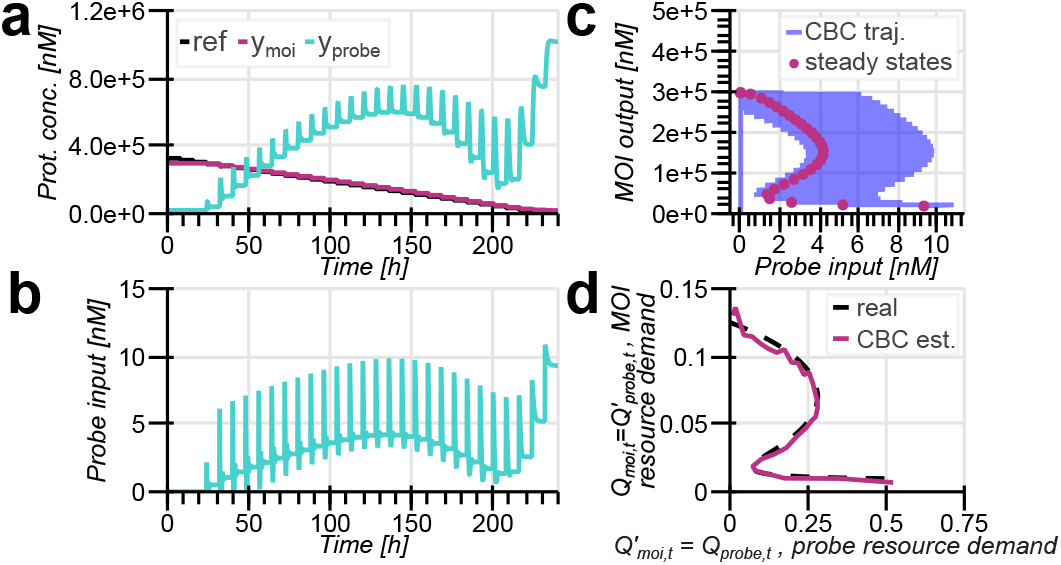
Simulation of MOI characterisation with CBC. (a–b) CBC trajectory with proportional control *u* = *K*_*p*_(*y*_*moi*_ *−R*) [7]. Transient spikes in *u* and *y*_*probe*_ correspond to switching to a new reference, which sharply increases the error, and decay as the new steady state is approached. (c) CBC bifurcation curve for the probe’s steady-state input vs the output of the MOI competing with it for resources. (d) Dashed: resource demand bifurcation curve *C*_*moi*_ relating the steady-state resource demands experienced and exerted by the characterised MOI, retrieved analytically according to (11). Solid: *C*_*moi*_ estimated from experimental data by applying the 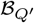 and ℬ_*Q*_ mappings.

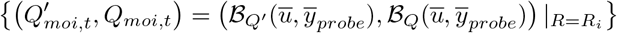

relate the resource demands experienced and exerted by the module in the steady state. Fig. 4d shows that the resultant estimates closely follow the real resource demand bifurcation curve, determined by analytically solving 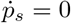 to obtain:

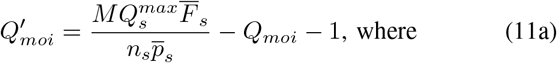

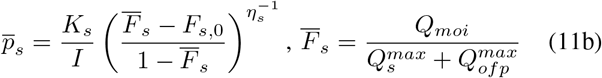

### C. Prediction of resource competition between modules

How can one predict the steady-state behaviour of two modules *moi*_1_ and *moi*_2_ hosted by the same cell, provided that they have been characterised with CBC as per Section III-B?

Recall that the resource demand exerted by one module is experienced by the other (and vice versa), so 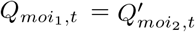 and 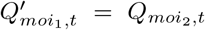. Hence, the two resource demand bifurcation curves 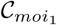 and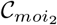 can be plotted on the same axes. For the overall system to be in steady state, both modules must be in their respective equilibria given by their bifurcation curves. The possible steady states of the combined gene expression system are thus given by intersections between 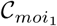 and 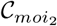. By mapping these intersections from resource demands back onto MOI outputs, the modules’ possible output combinations can be retrieved. Notably, neither the bifurcation curve’s retrieval nor the output prediction step assume any prior knowledge of the MOIs’ architecture or properties and solely rely on experimental data.

Our method’s predictive power is confirmed by simulation in Figure 5, where trajectories of two self-activating genetic switches converge from different initial conditions towards the possible output combinations corresponding to intersections between resource demand bifurcation curves. Importantly, our protocol may not merely estimate the steady-state outputs, but also allow one to identify qualitative changes in a system’s steady-state performance. For instance, repeating the procedure for a switch with greater chemical induction produces a shifted resource demand bifurcation curve 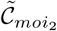 which has fewer intersections with 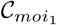 in Fig. 5c. This is consistent with the system having fewer possible steady-state outputs as shown in Fig. 5d.

**Fig. 5.**
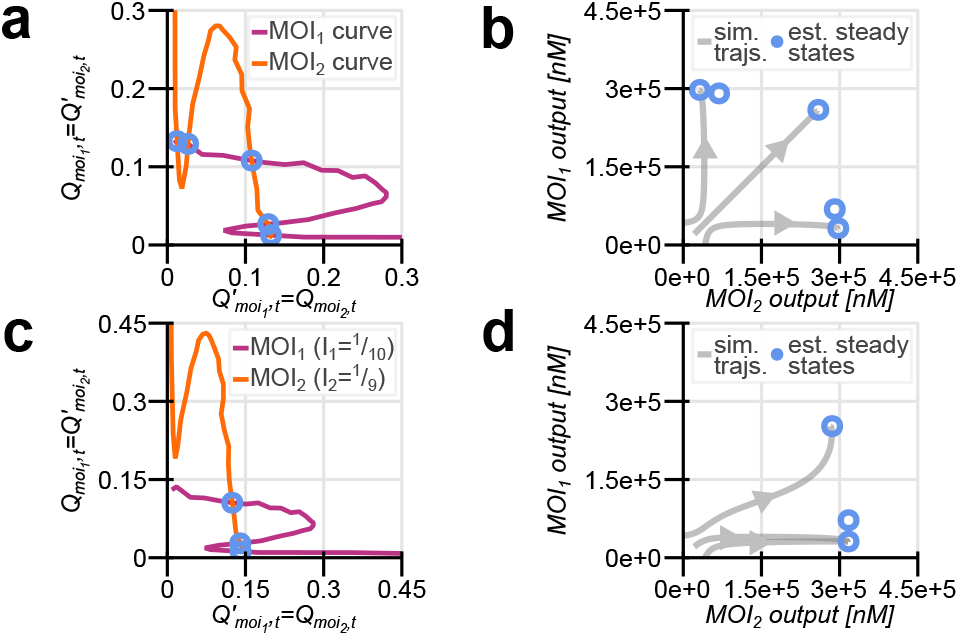
CBC characterisation predicts steady states of identical self-activating genetic switch modules competing for the cell’s resources. (a) Intersections between the superimposed estimated resource demand curves 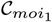 and 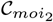 reveal possible steady states of the overall system (circles). (b) Circles: module outputs corresponding to the identified system steady states. Lines: simulated trajectories of the two-switch systems converging close to the identified steady states’ estimated outputs. (c–d) Steady-state prediction repeated for the case when the second switch is induced with *I* = 1/9 as opposed to *I* = 1/10 for the first switch here and both switches in a–b.

## IV. PROTOCOL EXTENSION TO A COMPLEX RESOURCE-AWARE CELL MODEL

Our characterisation protocol in Section III-B is based on CBC, whose model-free nature makes it applicable to systems with potentially unknown dynamics [7], [8], [13]. Likewise, in Section III-A knowledge of the probe module’s dynamics is not required, as the mappings 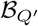 and ℬ_*Q*_ capturing the probe module’s resource competition properties are reconstructed simply by interpolation between experimentally acquired data points. Although translation of probe inputs and outputs into resource demands in Section III-A does assume a Hill dependence of protein expression rates on resource competition captured by the demands {*Q*_*i*_}, this relation is not arbitrary. Rather, it is mechanistically explained by competitive gene expression machinery binding dynamics and represents an empirical observation reproduced by different resource-aware models with widely ranging starting assumptions [1], [2], [5], [6]. Our method can thus be considered model-agnostic as long as the tested system’s properties are consistent with this Hill dependence that is commonly observed across different experiments and even model organisms [1], [2], [4]–[6].

To illustrate this, in Fig. 6 we simulate our protocol using a different, more complex cell model than the one in Sections II– III. We adopt a coarse-grained resource-aware mechanistic cell model from [1], originally implemented in Python for [16] using the JAX library for efficient computation [17]. This framework explicitly considers different steps of gene expression (rather than treating it as a single reaction in (1)) and captures dynamic regulation of ribosome synthesis and nutrient metabolism [1]. Despite the model’s greater complexity, our method still makes effective predictions in Fig. 6, supporting the promise of *in vivo* applicability to more complex real biological systems which may involve unknown interdependencies between our measured and actuated parameters.

**Fig. 6.**
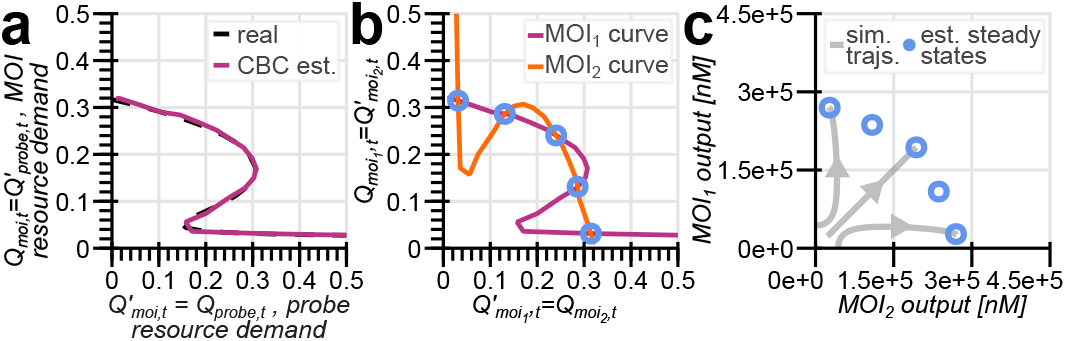
Simulation of our protocol with a coarse-grained mechanistic resource-aware cell model, for the example of a self-activating genetic switch module. (a) Real resource demand bifurcation curve *C*_*moi*_ (dashed; retrieved analytically as per [12]) and its estimate obtained by CBC characterisation (solid). (b) Intersections between estimated resource demand curves 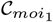 and 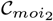 reveal possible steady states (circles) of two identical switches competing for the cell’s resources. (c) Steady-state outputs of the two system predicted by our method (circles) and simulated trajectories of such a two-switch system converging to the identified steady states (lines).

## V. CONCLUSION

We have described and simulated a novel experimental protocol which leverages automatic feedback control to characterise a synthetic gene circuit module’s resource competition properties. Designed to provide a more complete view of module behaviour than extant open-loop experimental approaches, our method reliably predicts steady-state behaviours of two genetic modules competing for a cell’s shared resources.

Our work contributes to a growing body of methods that aim to overcome the limitations of conventional experimental techniques by combining microscopy, real-time single-cell control, and computational methods in order to optimise the experimental design for maximum informativeness [10], [18] and observe difficult-to-capture cell behaviours [7], [9]. Hence, our approach is well-positioned to benefit from the emerging tools in synthetic biology, automated culturing, and control algorithms developed for these novel experimental methods. This may help to tackle the potential hurdles to our protocol’s practical implementation and wide adoption, such as long experimental timescales (e.g. bifurcation curve acquisition takes 240 *h* in our case, characteristically for CBC in biology [7]) and the need for robustness to intercellular variability and noisy measurements and gene expression dynamics.

Our method’s flexibility and model-agnosticism allow it to incorporate and combine the advantages of different experimental tools. A wide range of control strategies can be used to calculate feedback signals for CBC. More advanced solutions like model-predictive control, rather than the simple proportional feedback controller deployed here, can accelerate data collection and improve robustness to noise [7]. Optimal experimental design can ensure that references chosen for CBC tracking are maximally informative and fully cover the bifurcation curve [18]. Meanwhile, the automated cell culture platform suggested for CBC experiments – and validated *in silico* using agent-based simulations – is a microfluidic chip with individual bacterial cells at the end of each of thousands of trenches [7]. In this setup, control signals can be applied individually to each cell, e.g. using light to steer an optogenetically controlled probe module instead of chemical inputs affecting all bacteria growing in the medium [10]. Simultaneously acquiring thousands of CBC trajectories, steered by control signals computed in an efficient and parallelised manner [10], [16], [17], can enable reliable population-average readings despite measurement noise and individual trajectories’ stochasticity, as well as accelerate data acquisition, e.g. by using different subsets of cells to capture different sections of the bifurcation curve. Focussing on genetic modules’ general resource competition properties, rather than a particular system’s characteristic, our method allows to choose any CBC probe module as long as its input-output responses are reflective of resource competition in the cell. Even if (unlike for chemically controlled probes in the Appendix) this property may not always be theoretically proven, it may still be established experimentally by testing different optogenetic systems [19].

CBC’s greater time- and resource-intensity compared to simpler setups is offset by the reusability of its results. A once-established probe can be used to test many modules – including new alternative probes, bypassing characterisation with constitutive reporters in Section III-A. Once a resource demand bifurcation diagram for a module is obtained, it can be used to make predictions about its behaviour alongside any other similarly characterised module. As CBC testing is performed for more modules, its usefulness will thus scale combinatorially, which could facilitate the designing of reliable modular genetic circuits acting as biomolecular sensors or memory devices, where input-output responses must be known despite resource availability fluctuations [11]. Another potential application is cell fate reprogramming, in which competition between genetic regulators is an important factor [3]. Resource-dependent bifurcations, which our analysis helps establish, have also been leveraged in biomolecular controllers increasing the evolutionary stability of biotechnologies [12].

## APPENDIX

We assume no knowledge of the module of interest’s properties, so all resource competition interactions in the cell must be estimated from observing the probe module described in Section II-B. The probe’s inputs {*u*} are known as they are calculated and recorded on a computer, and fluorescence outputs 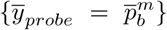 are measured with a microscope (Fig. 1b). *Theorem 1*, which follows from *Lemma 1*, proves that these values allow to unambiguously determine the resource demands of both the probe and the module co-expressed with it in the cell (i.e. the module of interest in our experiment).

### Lemma 1

For a given input 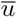 and resource demand 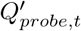 experienced by the probe module, exactly one steady-state output gene regulation function value 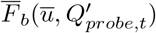 is possible. Moreover, for a given 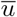 there are no such two values 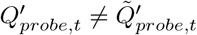 that 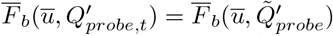.

*Proof:* Recalling that 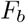 is a function of *u* and *p*_*ta*_ as per (4), let us express *p*_*ta*_ in terms of the steady-state 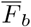and 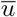, substitute it into 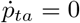, and rearrange the terms to get (12):

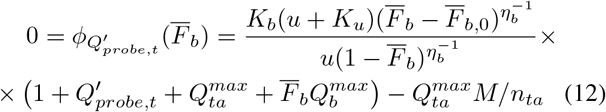

According to (4), *F*_*b*_ only takes values between *F*_*b*,0_ and 1. Since all parameters for biochemical systems like ours are non-negative, we note that 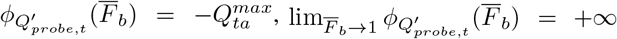, and 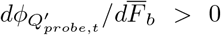. Rising monotonically from a negative to a positive infinite value, the continuous function 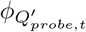 must therefore have exactly one root for which 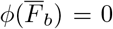. Then, let us observe that for 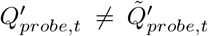 and any 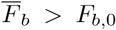, we have 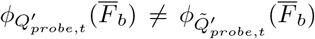 Hence, the roots of the two functions must be different. □

### Theorem 1

The steady-state resource demands experienced by the probe due to competing genetic modules, as well as the demand exerted by the probe, can be unambiguously determined from its input and output. That is, there exist mappings 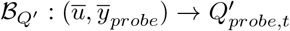 and 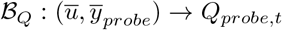 which become bijections between 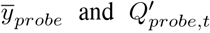 or between 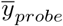 and 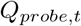 respectively, if 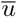 is fixed.

*Proof*. Set 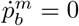 and 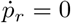then recall (2) to get:

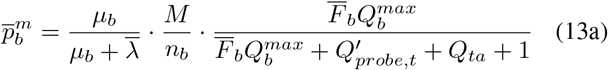

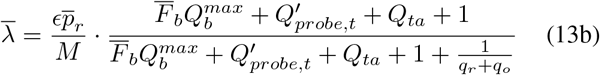

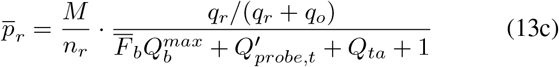

According to (13), for 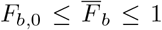 we have 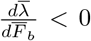 and thus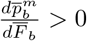. Consequently, 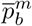 rises monotonically from the lowest-possible to the greatest-possible steady-state output value as 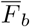 increases. Hence, there is a one-to-one match between the output gene’s steady-state regulatory function and its measured mature protein concentration. Combined with *Lemma 1*, this means that for a given input 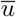, there can be no such two values 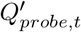and 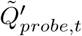 for which 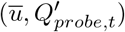 and 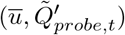 give rise to the same probe output 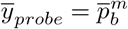, so 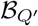 is a bijection. Then, due to *Lemma 1* the resource demand exerted by the probe is matched to the input-output pair via (14), making *ℬ*_*Q*_ a bijection as well. □

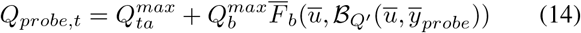

## ACKNOWLEDGMENT

We thank Sara Maria Brancato, Prof. Lucia Marucci and Dr Ludovic Renson for insightful discussions.

## Notes

This work was supported in part by the Engineering and Physical Sciences Research Council (EPSRC) grant EP/Y014073/1.

https://github.com/KSechkar/CoBa_CRAB

### Competing Interest Statement

The authors have declared no competing interest.

### Summary of Updates

Text rewritten for clairty; figures improved; typographical errors fixed.

